# Local delivery of SBRT and IL-12 to Murine PDAC Tumors Modulates Hematopoiesis

**DOI:** 10.1101/2025.01.10.632406

**Authors:** Tara G. Vrooman, Emily R. Quarato, Noah A. Salama, Maggie L. Lesch, Angela L. Hughson, Yuko Kawano, Gary Hannon, Sidney Lesser, Jian Ye, Sarah L. Eckl, Edith M. Lord, David C. Linehan, Nadia Luheshi, Haoming Qiu, Jim Eyles, Laura M. Calvi, Scott A. Gerber

## Abstract

**Background:** Standard of care therapies such as radiotherapy and chemotherapy have shown little efficacy against pancreatic ductal adenocarcinoma (PDAC). Immunotherapy is a newly emerging form of treatment that has shown promise; however, toxic systemic effects resulted in limited use in the clinic. Shifting from systemic to local delivery of cancer therapeutics reduces adverse systemic effects and increases response rates in multiple malignancies. Importantly, the effects of tumor-targeted therapies on distal tissues, such as the bone marrow, have not been thoroughly investigated.

**Methods:** Using a murine model of PDAC, we treated tumors with targeted stereotactic body radiation therapy (SBRT) and intratumoral interleukin-12 (IL-12). 13 days-13 months after tumor injection, the cells in the tumor, blood, and bone marrow were analyzed for therapy-induced changes. Hematopoietic cell numbers and lymphocytes were quantified by flow cytometry, and cytokine levels were quantified by enzyme-linked immunosorbent assays (ELISAs).

**Results:** We demonstrated that although SBRT/IL-12 delivered locally to PDAC tumors successfully eradicated primary disease, it also induced significant acute and long-term effects in the bone marrow. Within days of intratumoral SBRT/IL-12 treatment, we observed acute lymphopenia in the blood, accompanied by an immunostimulatory response in the bone marrow characterized by an increase in hematopoiesis. Long-term effects included a decrease in hematopoietic stem cells (HSCs) and skewing toward a myeloid lineage bias, which could indicate premature aging of the HSC population.

**Conclusions:** These findings demonstrate that despite being locally delivered to the tumor, SBRT/IL-12 therapy exerts significant effects on the distal bone marrow, reinforcing the need for further investigations into the long-term systemic immunological outcomes of localized cancer treatments.

**Key Messages:** <u>What is already known on this topic</u>: Systemic cancer therapies used to combat pancreatic ductal adenocarcinoma (PDAC) often induce toxic systemic effects. Local delivery of radiation and immunotherapy reduces adverse effects; however, the systemic spread of these therapies and the resulting effects on distal tissues such as the bone marrow have yet to be elucidated.

<u>What this study adds</u>: Intratumoral delivery of stereotactic body radiation therapy (SBRT) and interleukin-12 (IL-12) augment hematopoiesis in the bone marrow soon after treatment and induce long-term alterations in the hematopoietic stem cells (HSCs). These effects are mainly a result of IL-12 that is transiently increased in the bone marrow after treatment.

<u>How this study might affect research, practice, or policy</u>: Targeted SBRT/IL-12 therapy induces long-term systemic effects on the bone marrow, indicating the need for further investigation of the systemic spread of locally delivered therapeutics.

## 1. Background

Immunotherapy is an emerging cancer treatment that harnesses the body’s immune system to recognize and destroy cancer cells, offering a promising alternative to traditional cytotoxic therapies like chemotherapy and radiation. Systemically administered immunotherapies have transformed cancer treatment, particularly through durable responses seen with immune checkpoint inhibitors (ICIs) in various advanced malignancies [1, 2]. However, many patients do not respond to these therapies due to multiple tumor resistance mechanisms [3]. Additionally, systemic immunotherapy is hampered by dose-limiting toxicities, restraining treatment efficacy [4]. Localized delivery can circumvent these barriers, allowing for higher concentrations of the therapeutic agent at the tumor site while also minimizing toxicity [5]. Interleukin-12 (IL-12) is a prime example of an immunotherapeutic that has shifted from systemic to localized delivery due to its significant toxicity when administered intravenously. IL-12, a pleiotropic antitumor cytokine capable of activating both innate and adaptive immunity, exhibited significant toxicity including fatigue, dyspnea, leukopenia, and even life-threatening complications when administered systemically in phase II clinical trials [4, 6]. Numerous delivery systems have since been developed to administer IL-12 intratumorally, including lipid nanoparticles, microspheres, genetically engineered cells, and viral gene therapy [7]. However, it is important to note that while these therapies reduce systemic side effects, they do not eliminate them entirely. Moreover, there are few studies that thoroughly examine the systemic effects of locally-administered therapies.

Our laboratory has developed a combination therapy incorporating targeted stereotactic body radiotherapy (SBRT) to the pancreatic tumor followed by local administration of IL-12 using various nanoparticle delivery platforms [8, 9]. This localized dual approach has resulted in the elimination of murine primary and metastatic tumors through robust anti-tumor immune responses [8, 9], prompting clinical evaluation of this treatment (NCT06217666). However, to fully appreciate the potential of this combination therapy, it is vital to understand the effects of IL-12 and SBRT both locally and distally. Systemic administration of IL-12 has been shown to induce cytokine storms characterized by the overproduction of pro-inflammatory cytokines such as interferon-γ (IFNγ), tumor necrosis factor-α (TNFα), and interleukin-6 (IL-6) [10]. Additionally, systemic IL-12 can reduce lymphocytes, monocytes, and neutrophils in the blood, triggering emergency hematopoiesis in the bone marrow [11]. Likewise, SBRT, even when the bone compartment is not targeted, can alter the complex cellular network in the bone marrow, leading to long-term changes in hematopoiesis and myelopoiesis, potentially resulting in premature aging of the bone marrow [12–15]. Given the known systemic effects generated by each of these as a monotherapy, it is essential to investigate the combined systemic impact of locally delivered IL-12 and SBRT to fully understand the potential effects of this treatment and maximize patient safety.

Here we report that local administration of SBRT and IL-12 eradicated primary disease, with IL-12 concentrations highest in the tumor as expected. Detectable, but significantly lower levels, were also identified in the blood and bone marrow. Consequently, we observed acute lymphopenia and granulocytosis in the peripheral blood, as indicated by complete blood counts (CBC). These hematological changes coincided with significant reductions in bone marrow hematopoietic stem cell (HSC) populations, most notably 13 days post-therapy. Although the effects on the bone marrow were less pronounced, they persisted even 16 months later. Notably, most of these changes were attributed to IL-12 rather than SBRT, indicating the cytokine was the driver of the observed responses. These findings demonstrate that local SBRT and IL-12 therapy is capable of producing systemic changes, particularly on bone marrow progenitor cells. However, these effects are minimal, comparable to that of conventional cytotoxic therapies used in the clinic [16], and likely acceptable given the positive risk-benefit ratio of the treatment for pancreatic cancer.

## 2. Materials and Methods

### 2.1 Cell Line

KP2 cells, derived from KPC mice, were provided by Dr. David DeNardo [17]. These cells were engineered to stably express firefly luciferase (KP2.1-luc) and were grown using MAT/P media (US patent No. 4.816.401) supplemented with 5% fetal bovine serum (Hyclone) and 1% penicillin/streptomycin (Gibco) and incubated at 37°C at 5% CO₂.

### 2.2 *In vivo* Animal Studies

Six- to eight-week old female C57BL/6J mice from The Jackson Laboratory were used for all experiments. All mice were kept on a 12-hour light/dark cycle with ventilated cages, bedding and nesting material, and enrichment houses. All experiments used mice randomized into respective groups with blinding introduced throughout studies. All experiments were approved by the University of Rochester Committee on Animal Resources and performed in compliance with the NIH, ARRIVE [18] and University of Rochester guidelines for the care and use of animals.

### 2.3 Orthotopic Tumor Implantation

Mice were injected with buprenorphine (Fidelis Animal Health) and anesthetized using vaporized isoflurane (Vet One) and a 10-mm laparotomy incision was made to expose the pancreas. 25,000 KP2.1-luciferase expressing cells in PBS (Gibco) were mixed in a 1:1 dilution with Matrigel (Corning) and injected in a volume of 40µL into the tail of the pancreas. Two titanium fiducial clips (4-mm, Horizon) were inserted on either side of the tumor to provide targeting during radiation. Mice were monitored daily for tumor burden.

### 2.4 Radiation Therapy

Six days after surgery, tumor-bearing mice were anesthetized with isoflurane and received 6 Gy SBRT for 4 consecutive days (days 6-9 post-tumor injection) using a 5-mm collimator on a Small Animal Radiation Research Platform (SARRP, XStrahl). Tumors were targeted with CT guidance, and treatments were administered using Muriplan software as previously described [8, 9].

### 2.5 IL-12mRNA Therapy

IL-12mRNA and scrambled mRNA (scRNA) in lipid nanoparticles were provided by AstraZeneca [19]. Twenty-four hours after the final SBRT fraction, anesthetized tumor-bearing mice underwent a 10-mm laparotomy and IL-12mRNA or scRNA lipid nanoparticles (0.5µg in 25µL) were directly injected into the tumor using an 18-gauge syringe.

### 2.6 Bioluminescent Imaging

*In vivo* tumor growth was monitored using an IVIS Spectrum *In Vivo* Imaging System (IVIS, PerkinElmer) where anesthetized mice were injected with D-luciferin (2.5mg/100 µL in PBS, Invitrogen) and peak bioluminescence (p/sec/cm^2^/sr) was calculated within a tumor regions of interest (ROIs).

### 2.7 Antibody Depletion

Mice cured of KP2.1-luc tumor cells following SBRT/IL-12 therapy were injected subcutaneously with an antibody cocktail (Bio X Cell) containing 200µg each of rat anti-mouse CD4 (IgG2a, GK1.5), rat anti-mouse CD8 (IgG2a, 53-6.7), and rat anti-mouse NK1.1 (IgG2a, PK136) in 100µL PBS or rat isotype control (IgG2a, C1.18.4) every 3 days for 30 days. Tumor burden was monitored by IVIS.

### 2.8 Blood Chemistry Analysis

Blood was collected from the retroorbital sinus and serum isolated. Serum was sent out for testing at VRL Animal Health Diagnostics for amylase and lipase levels.

### 2.9 qPCR Assay

The pancreases from cured and age-matched naïve mice were digested using 30% collagenase in HBSS (30 minutes, 37°C, Sigma-Aldrich). Homogenates were prepared for gDNA isolation using a DNeasy Blood and Tissue Kit from Qiagen. Luciferase expression was assayed using iTaq Universal SYBR Green Supermix (BioRad) and a CFX96 Real-Time System (C1000 Thermocycler, BioRad).

### 2.10 Peripheral Blood Collection and Complete Blood Count (CBC) Analysis

Blood was collected from mice via the submandibular vein into a BD Microtainer (BD). 50µL of whole blood was assayed on the HESKA HT5+ for CBCs.

### 2.11 Isolation of Bone Marrow and BMEF Collection

Following euthanasia, bilateral tibiae, femora, and pelvic bones were collected, cleaned of soft tissues, and cut into 3-4 pieces. Bone pieces were crushed using a mortar and pestle to release bone marrow into 3ml of PBS. Bone marrow was passed through an 18G needle to disassociate clumps and pelleted by centrifugation at 1200rpm for 5 minutes. The bone marrow extracellular fluid (BMEF) was collected for protein analysis. Red blood cells were lysed (156mM NH_4_Cl, 127μM EDTA, and 12mM NaHC_3_) for 5 minutes at room temperature and filtered through a 100μm filter. Cell numbers were determined using the TC20 (Biorad) automated cell counter using trypan blue (Sigma-Aldrich) to exclude dead cells.

### 2.12 Flow Cytometry

Bone marrow cells were processed as described above. Up to 10x10^6^ cells were incubated for 30 minutes at 4°C in 100µL of PBS and 2% FBS containing fluorophore-conjugated antibodies and 2µg/mL DAPI (Invitrogen) to label dead cells. The following conjugated antibodies were used for staining: CD34-FITC (RAM34), CD48-PE-Cy7 (HM48-1), CD45-APC-Cy7 (30-F11), c-kit-APC (2B8), Ter119-PerCP-Cy5.5 (Ter-119), B220-PerCP-Cy5.5 (RA3-6B2), CD3e-PerCP-Cy5.5 (145-2C11), CD45-PerCP-Cy5.5 (30-F11), CD11b-AF700 (M1/70), CD4-PE-Cy5 (RM4-5), CD62L-PE-Cy7 (MEL-14), and Gr1-PerCP-Cy5.5 (RB6-8C5) all from BD Pharmingen, CD16/32-APC-Cy7 (517011E), IgM-FITC (RMM-1), CXCR3-BV650 (CXCR3-173), F4/80-FITC (BM8), and CD206-APC (C068C2) from Biolegend, Sca-1-BV650 (D7), MHCII(I-A/I-E)-BV786 (M5-114), and CD41-BUV395 (MWReg30) from BD Optibuild, CD19-PE-CF594 (1D3), CD8α-BUV395 (53-6.7), CD3ε-BUV737 (145-2C11), Ly6C-BV605 (AL-21), CXCR4-PE-CF594 (2B11/CXCR4), and Ly6G-BUV395 (1A8) from BD Horizon, Flt3-PE-Cy5 (A2F10, Invitrogen), and B220-APC (RA3-6B2, Tonbo). All samples were run on an LSRFortessa flow cytometer (BD Biosciences) and analyzed using FlowJo version 10.8 (Tree Star).

### 2.13 ELISA Assays

The tumor/pancreas, bone marrow extracellular fluid, and blood serum were collected and tumors/pancreases homogenized for 1 hour in cell lysis buffer #2 (R&D Systems) with protease inhibitors on ice. Supernatants were assayed for IL-12 or IFNγ proteins using ELISA Max Deluxe kits (Biolegend) according to manufacturer’s instructions.

### 2.14 Statistical Analysis

GraphPad Prism 10 Software was used and P values of <0.05 were considered significant. Unless noted otherwise, all summary values represent the mean and standard deviation of individual data points. One-way ANOVAs or unpaired t-tests were used as appropriate for the data presented with Tukey multiple comparisons corrections as needed.

## 3. Results

### 3.1 Combination SBRT/IL-12 therapy eradicates PDAC tumors

Using a previously developed murine model of PDAC [8, 9], we directly treated the tumor with SBRT and IL-12mRNA or a scrambled RNA control (scRNA) (**Figure 1A**). In each experiment, mice were randomized into four different treatment groups: no radiation and scRNA hereafter referred to as Untreated (UT), SBRT and scRNA hereafter referred to as SBRT, no radiation and IL-12mRNA (IL-12), and SBRT/IL-12. Combination SBRT/IL-12 therapy reliably cured PDAC tumors, whereas all tumors persisted in SBRT only and UT mice, and IL-12 treatment alone failed to consistently eliminate tumors (**Figure 1B**). Additionally, when mice cured for over 100 days were depleted of T and NK cells, there was no tumor recurrence suggesting complete tumor eradication (**Supplemental Figure 1A and 1B left and middle**). Quantitative PCR analysis detecting luciferase expressed only by PDAC tumor cells confirmed the lack of dormant tumor cells in the pancreas 100 days after tumor cure (**Supplemental Figure 1B right**). To assess the risk of pancreatitis resulting from local therapy, blood chemistry analysis was performed on naïve mice and mice cured for 30 days and 143 days. Our analysis revealed normal amylase and lipase levels, recognized indicators of pancreatitis, in both naïve and previously cured mice (**Figure 1C**) [20]. These data suggest that SBRT/IL-12 therapy completely eradicates PDAC tumors and does not result in detrimental side effects such as pancreatitis.

**Figure 1:**
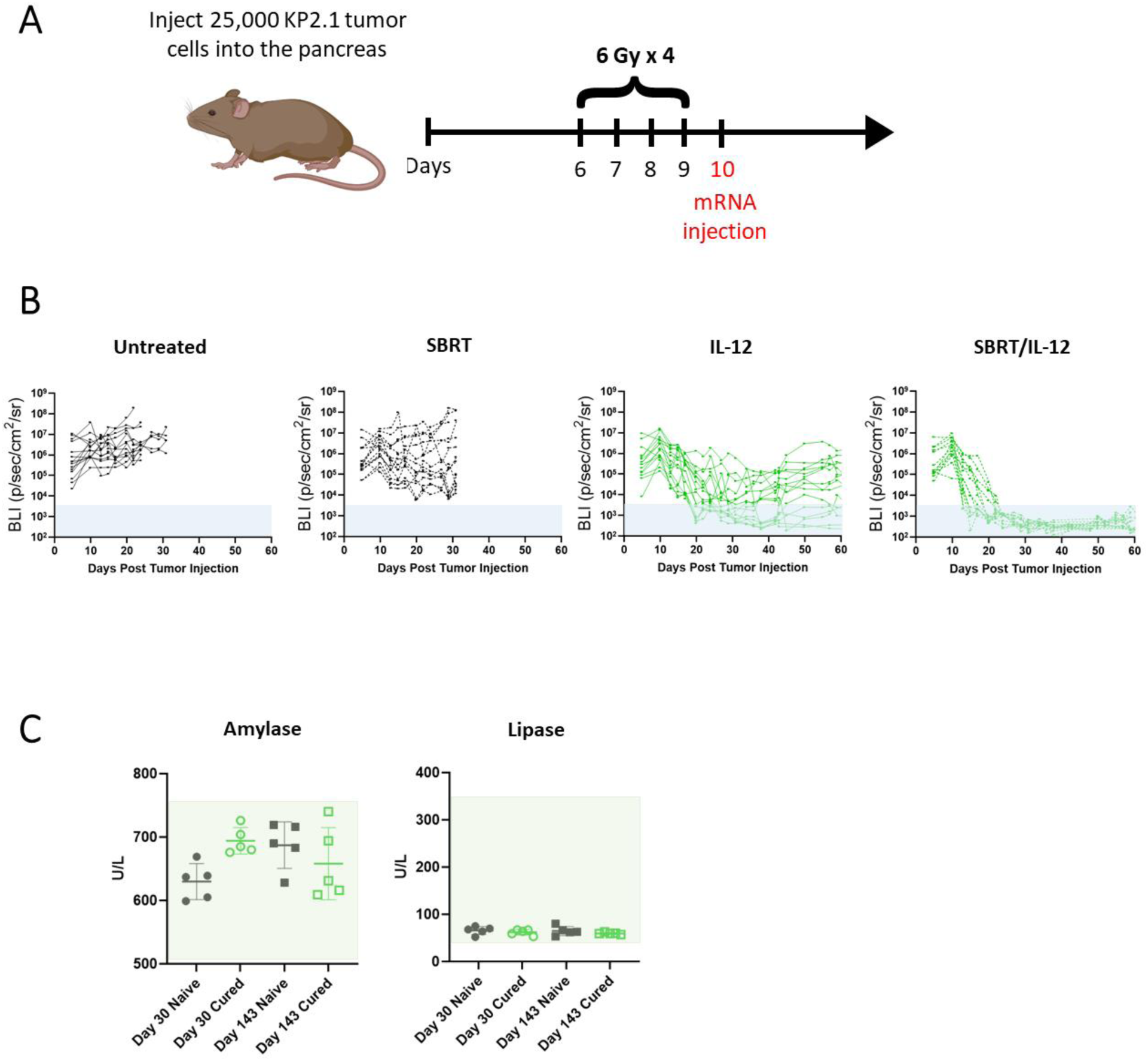
SBRT/IL-12 therapy cures pancreatic cancer in a murine model of PDAC. **A** Treatment timeline of tumor injection, SBRT, and IL-12 administration. **B** Tumor burden BLI in each treatment group. Shaded blue area represents a BLI measurement at baseline without tumor. **C** Blood chemistry testing of amylase and lipase levels in cured or age-matched naïve mice 30 days and 143 days after tumor injection. Shaded areas represent the normal range of values (512-744 for amylase, 47-356 for lipase). Untreated and SBRT-treated mice received scRNA as a control. n = 5-15.

### 3.2 Intratumoral IL-12 and IFNγ concentrations are increased 24 hours after IL-12mRNA injection

Given that IL-12mRNA is directly injected into the tumor in our model, we performed enzyme-linked immunosorbent assays (ELISAs) to quantify the concentration of translated intratumoral IL-12 protein and IFNγ, a common downstream mediator of IL-12 [21]. A significant increase of IL-12 protein was detected in the tumor 24 hours after IL-12mRNA injection that quickly dissipated 72 hours post injection (**Figure 2A**). Likewise, IFNγ was significantly increased in the tumor one day after IL-12 injection, and, in contrast to IL-12, remained elevated 72 hours later (13 days post tumor injection) (**Figure 2B**). These data confirm that IL-12mRNA results in an increase of intratumoral IL-12 protein and IFNγ 24 hours after treatment.

**Figure 2:**
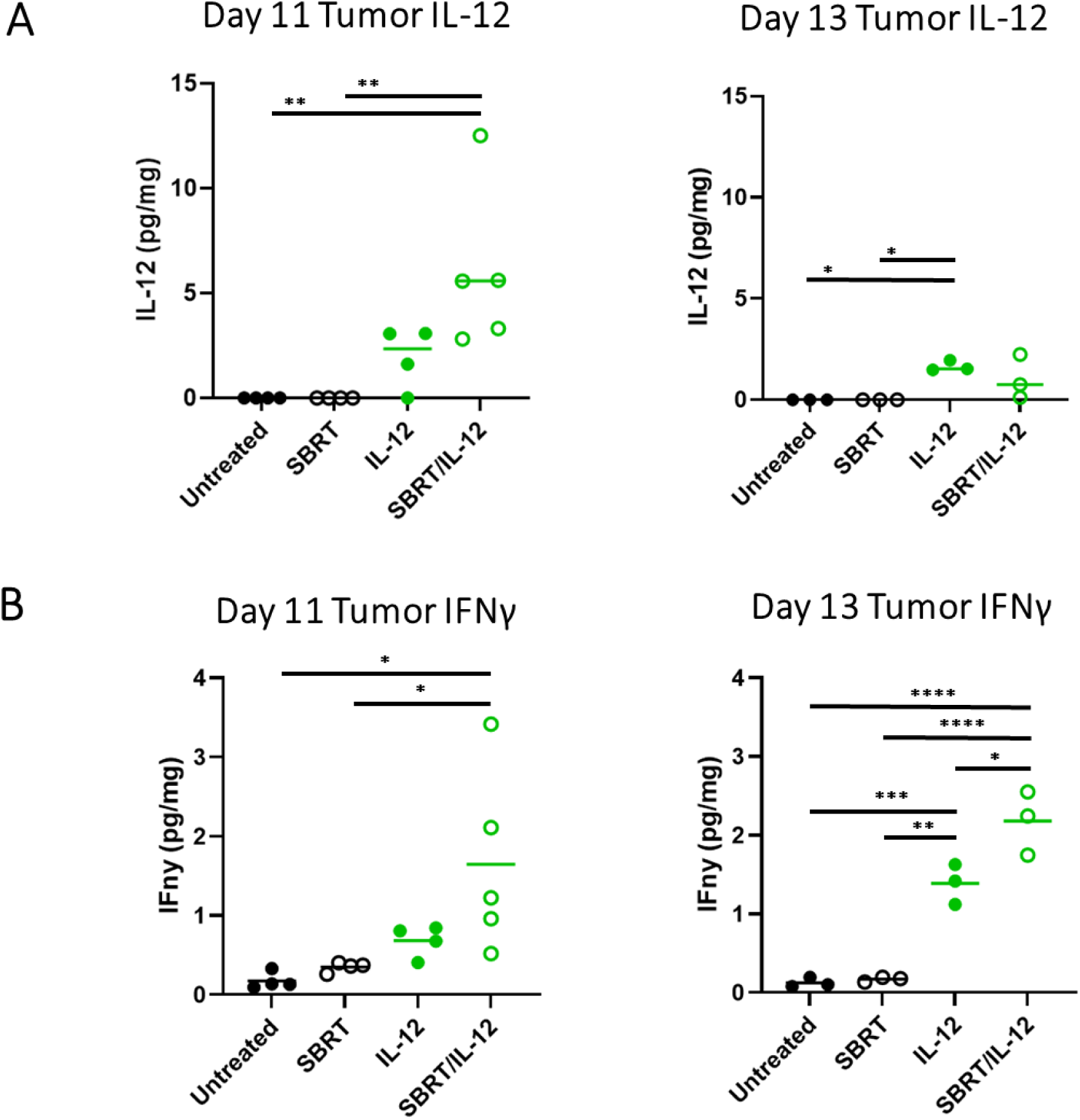
Intratumoral IL-12 and IFNγ are detectable 24 hours after IL-12 injection. ELISA assay results showing **A** IL-12 and **B** IFNγ protein levels in the tumor 24 hours and 72 hours after IL-12 injection (days 11 and 13 after tumor injection respectively). Untreated and SBRT-treated mice received scRNA as a control. n = 3-5; *p<0.05; **p<0.01; ***p<0.001; ****p<0.0001.

### 3.3 IL-12 treatment temporarily alters lymphocyte counts in the peripheral blood

ELISA analysis of the blood revealed elevations of IL-12 protein 24 hours after injection (**Figure 3A**). This effect was transient, as levels returned to baseline 72 hours after injection (day 13) (**Figure 3B**). To assess the effect of combination treatment on peripheral blood cells, we analyzed the CBCs for each treatment group. Lymphopenia was observed on day 13 after tumor injection in both groups treated with IL-12. These effects were not seen in the SBRT only or UT groups (**Figure 3C**). No significant differences were observed in monocyte and granulocyte densities (**Figure 3D-E**). Notably, IL-12 protein levels and all peripheral blood immune cells returned to normal levels one year after combination therapy, indicating this effect was transient (**Supplemental Figure 2**). These findings indicate temporary immune cell changes occur systemically after SBRT/IL-12 therapy.

**Figure 3:**
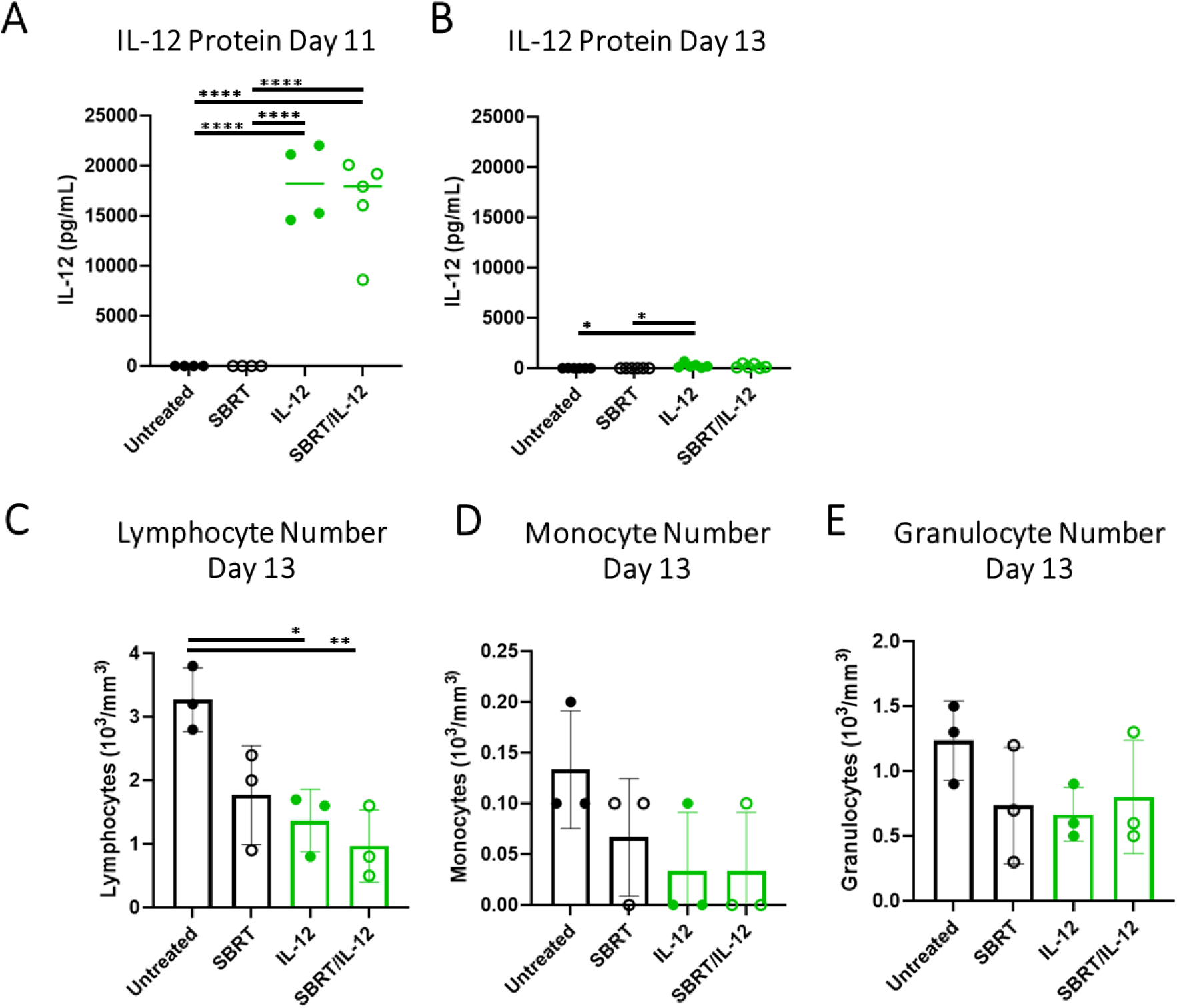
Lymphocyte numbers in the blood are altered after SBRT/IL-12 therapy. ELISAs on blood serum were performed **A** 24 hours (day 11) and **B** 72 hours (day 13) after IL-12mRNA (IL-12 and SBRT/IL-12) or scRNA (Untreated and SBRT) injection. Complete blood count (CBC) data revealed the number of **C** lymphocytes, **D** monocytes, and **E** granulocytes within the blood 13 days after tumor injection. n = 3; *p<0.05; **p<0.01.

### 3.4 IL-12 is transiently increased in the bone marrow after SBRT/IL-12 therapy

Due to changes in peripheral blood immune cells, we postulated that the bone marrow, the primary site of hematopoiesis, may be impacted in response to our treatment. We investigated whether IL-12 was detectable in the bone marrow even though IL-12mRNA was administered locally to the pancreatic tumor. IL-12 and IFNγ protein concentrations were assessed using ELISAs on the bone marrow extracellular fluid at four different timepoints (day 11, day 13, day 41, and 13-16 months post tumor injection). On day 11, a significant amount of IL-12 was detected in both groups that received intratumoral IL-12mRNA administration (**Figure 4A**). Of note, the concentration of IL-12 in the bone marrow was 30-fold less than the concentration in the tumor at the same time point. 13 days after tumor injection, IL-12 protein levels returned to normal and remained at baseline 13-16 months later (**Figure 4B-D**). Surprisingly, there were no significant differences in IFNγ expression between the SBRT/IL-12 treated group and the other treatment groups at any timepoint (**Figure 4A-D**). These data suggest that a significant concentration of IL-12 is present in the bone marrow 24 hours after IL-12mRNA injection into the primary tumor, but it quickly dissipates within 72 hours of injection. In contrast to our observations in the tumor, our findings indicate IL-12 within the bone marrow does not stimulate an increase in production of IFNγ in the combination treatment group at these timepoints.

**Figure 4:**
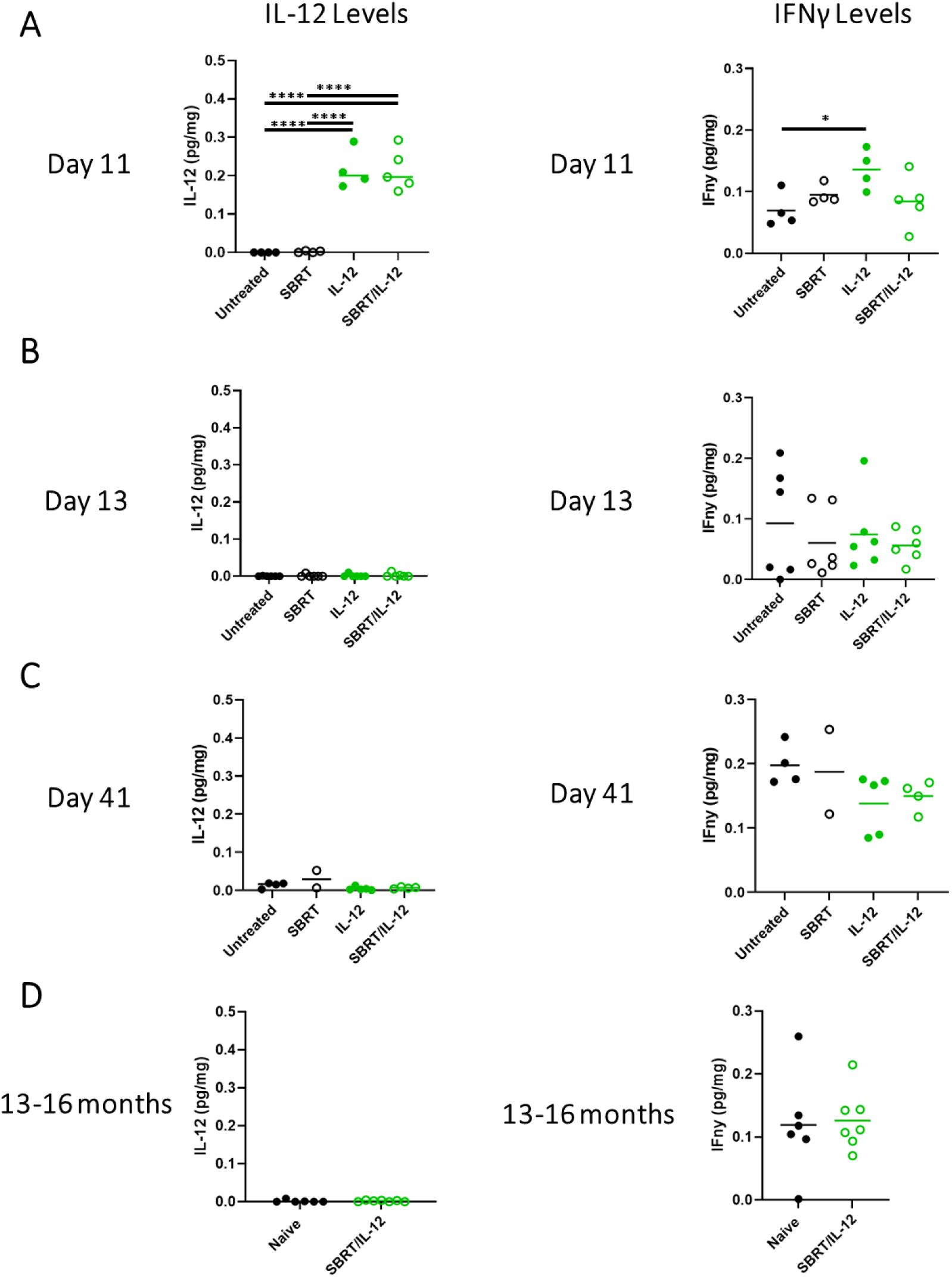
IL-12 and IFNγ levels are transiently increased in the bone marrow after treatment. ELISA assay results showing IL-12 and IFNγ protein levels in the tumor at **A** 11 days, **B** 13 days, **C** 41 days, and **D** 13-16 months after tumor injection. Untreated and SBRT-treated mice received scRNA as a control. n = 2-7; *p<0.05; ****p<0.0001.

### 3.5 SBRT/IL-12 therapy causes minor long-term changes in the bone marrow

Standard of care treatments such as radiotherapy and chemotherapy can modulate the cells of the bone marrow, causing a permanent decrease in the self-renewal capabilities of long-term hematopoietic stem cells (LT-HSCs) [15, 16, 22]. We investigated whether our therapy targeted to the pancreas has systemic effects on the bone marrow. To assess this, we performed flow cytometry using canonical markers to quantify the HSCs, progenitor cells, and their terminally differentiated populations within the bone marrow (**Supplementary Figure 3A & B**) [23]. The surface markers used to define each population are included in **Supplementary Table 1**.

These bone marrow cells include multiple subsets of HSCs, which produce all the cells in the lymphoid and myeloid lineages. For example, LT-HSCs possess *unlimited* self-renewal capabilities, and give rise to all erythroid, myeloid, and lymphoid cells, as well as platelets. Downstream of LT-HSCs, short-term HSCs (ST-HSCs), with *limited* self-renewal, differentiate into multipotent progenitor cells (MPPs) that further differentiate into myeloid and lymphoid lineages.

Specifically, MPP2 and MPP3 cells give rise to various myeloid progenitors including **1)** common myeloid progenitor cells (CMPs), **2)** granulocyte-monocyte progenitor cells (GMPs), **3)** megakaryocyte-erythroid progenitor cells (MEPs), and **4)** megakaryocyte progenitor cells (MKPs). MPP2 cells preferentially give rise to MEPs and MKPs, whereas MPP3 cells differentiate into GMPs that eventually become granulocytes and monocytes. On the lymphoid side, MPP4 cells, differentiate into common lymphoid progenitor cells (CLP) which can further mature into B cells, natural killer (NK) cells, or T cells (CD45+, CD3+).

We assessed each of these cell types after therapy and observed significant changes in the percentage of myeloid progenitors and stem cell progenitors after SBRT/IL-12 therapy (**Figure 5A**). Specifically, we discovered a significant decrease in LT-HSCs in the bone marrow 72 hours after IL-12 injection (day 13), which was sustained 13-16 months later (**Figure 5B**). This prolonged decrease in stem cells was mirrored by the ST-HSCs (**Figure 5D**). Notably, the percentage of LT-HSCs and ST-HSCs was unchanged between treatment groups by day 41 after tumor injection (**Supplemental Figure 4A**).

**Figure 5:**
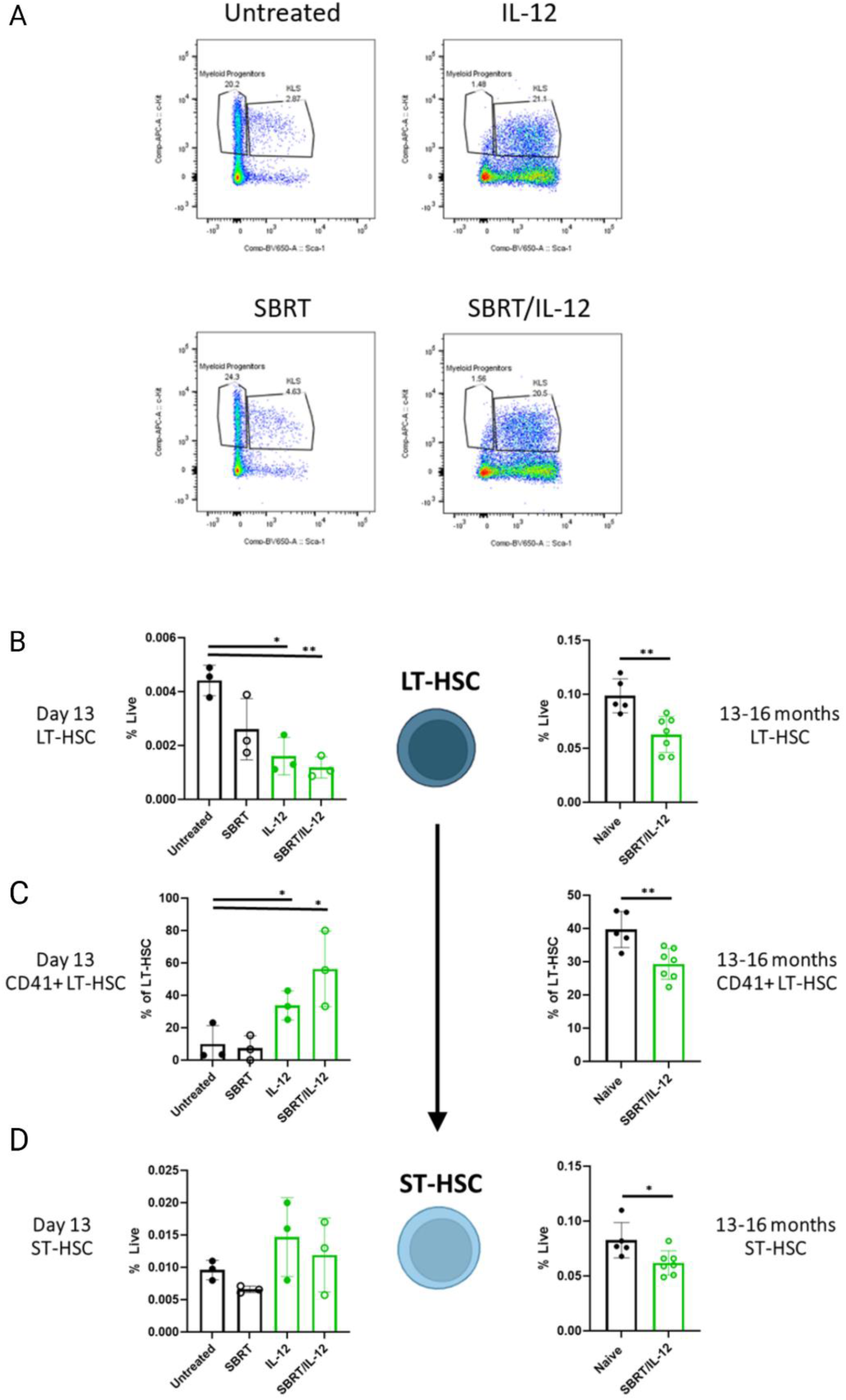
SBRT/IL-12 results in reduction of hematopoietic stem cell levels and skews cells towards myeloid differentiation. **A** Representative flow plots of myeloid progenitors and stem cell progenitor cells (KLS) in each of the four treatment groups (Untreated, SBRT, IL-12, and SBRT/IL-12). Quantification of **B** LT-HSCs, **C** CD41+ LT-HSCs, and **D** ST-HSCs within the bone marrow at day 13 and 13-16 months after tumor injection. Untreated and SBRT-treated mice received scRNA as a control. n = 3; *p<0.05; **p<0.01.

The phenotype of the LT-HSCs at day 13 after tumor injection revealed a significant increase in CD41 expression after SBRT/IL-12 therapy (**Figure 5C**). CD41 expression on stem cells indicates differentiation toward the myeloid lineage [24]. This was supported in our data by the increase in myeloid progenitor cells (MPP2 and MPP3) shortly after IL-12 therapy (**Figure 6A and Supplemental Figure 6A**). Although the MPP2 and MPP3 cells were increased, the downstream progenitors (CMPs, GMPs, MEPs, and MKPs) were decreased (**Figure 6A-C and Supplemental Figure 6B-C**). Similarly, the lymphocyte progenitor MPP4 cells were increased by combination therapy, which may be due to a need for additional T cells in response to the IL-12 (**Supplemental Figure 7**). Mature T cells within the bone marrow were unaffected by combination therapy (**Supplemental Figure 8**). These data demonstrate a significant activation of HSCs in response to localized therapy to pancreatic tumors.

**Figure 6:**
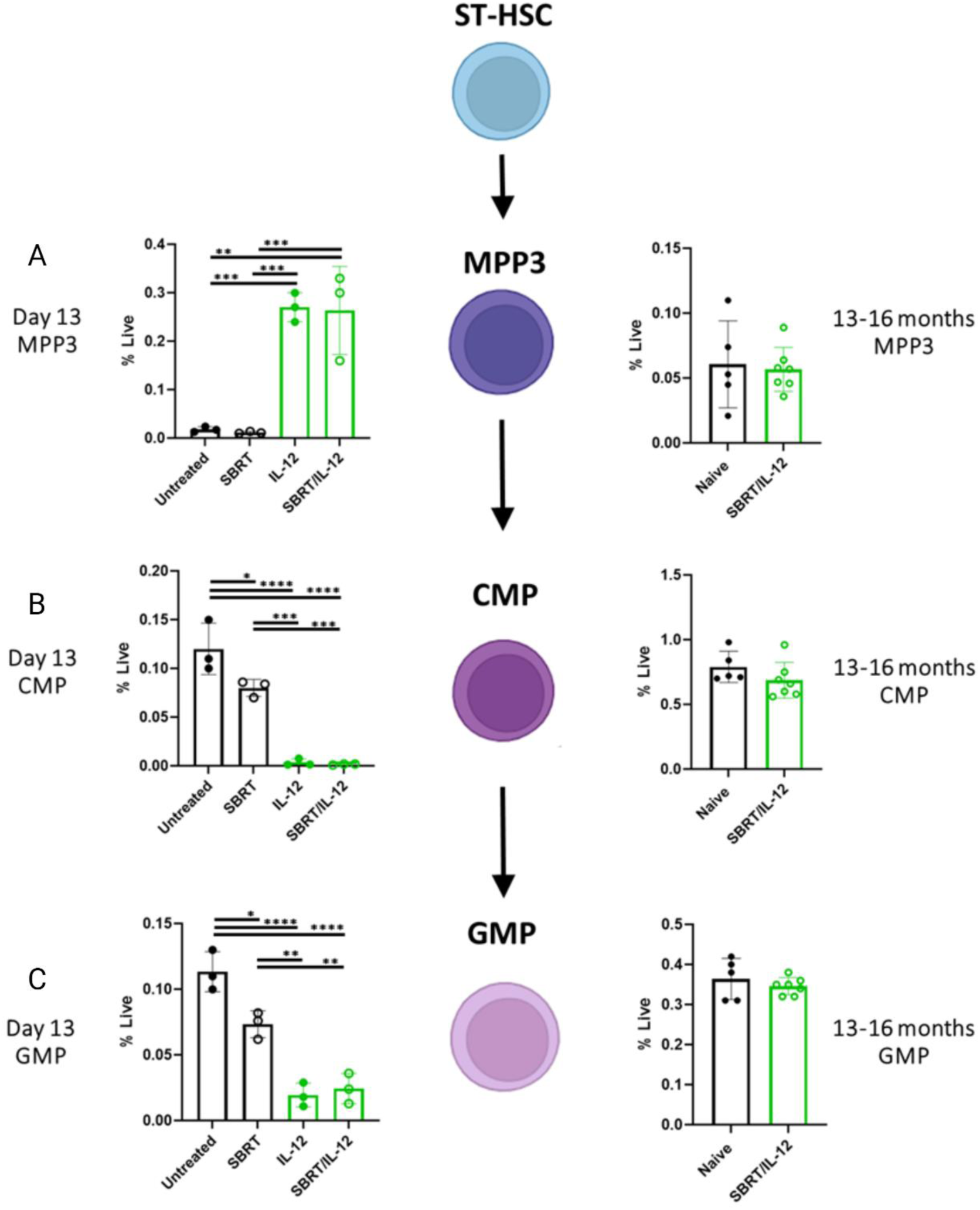
SBRT/IL-12 therapy alters the myeloid progenitors in the bone marrow. Quantification of **A** MPP3, **B** CMP, and **C** GMP cells in the bone marrow at day 13 and 13-16 months after tumor injection. Untreated and SBRT-treated mice received scRNA as a control. n = 3-7; *p<0.05; **p<0.01; ***p<0.001; ****p<0.0001.

### 3.6 IL-12 activates myeloid cells in the bone marrow

In the myeloid differentiation pathway, MPP3 cells differentiate into CMPs, then GMPs. Both cell types were significantly decreased in the bone marrow in the IL-12 and SBRT/IL-12 groups at day 13 (**Figure 6B & C**), but returned to normal levels by day 41 after tumor injection indicating a transient effect of therapy (**Supplemental Figures 4B & 5B**). As GMPs are progenitors for monocytes (CD45+, Ly6G-, Ly6C+) and macrophages (CD45+, Ly6G-, Ly6C-, F480+), we measured the production of terminally differentiated myeloid cells. We observed a significant increase in monocytes in the IL-12 treated groups only at day 13 (**Figure 7A**), whereas macrophage numbers were unchanged at day 13 (**Figure 7C**) but were elevated at day 41 (**Supplemental Figure 5B**). Both cell types exhibited an activation phenotype induced by IL-12 treatment, as evidenced by an increase in MHCII expression, a pro-inflammatory marker associated with T cell activation (**Figure 7B & D**) [25]. These differences were normalized by 13-16 months post treatment (**Supplemental Figure 5 and Figure 7B & D**), indicating there are no long-term effects in the myeloid cell compartments. Collectively, these data suggest that IL-12 initially induces activation and differentiation of monocytes and macrophages, however these changes are transient and return to baseline at later timepoints.

**Figure 7:**
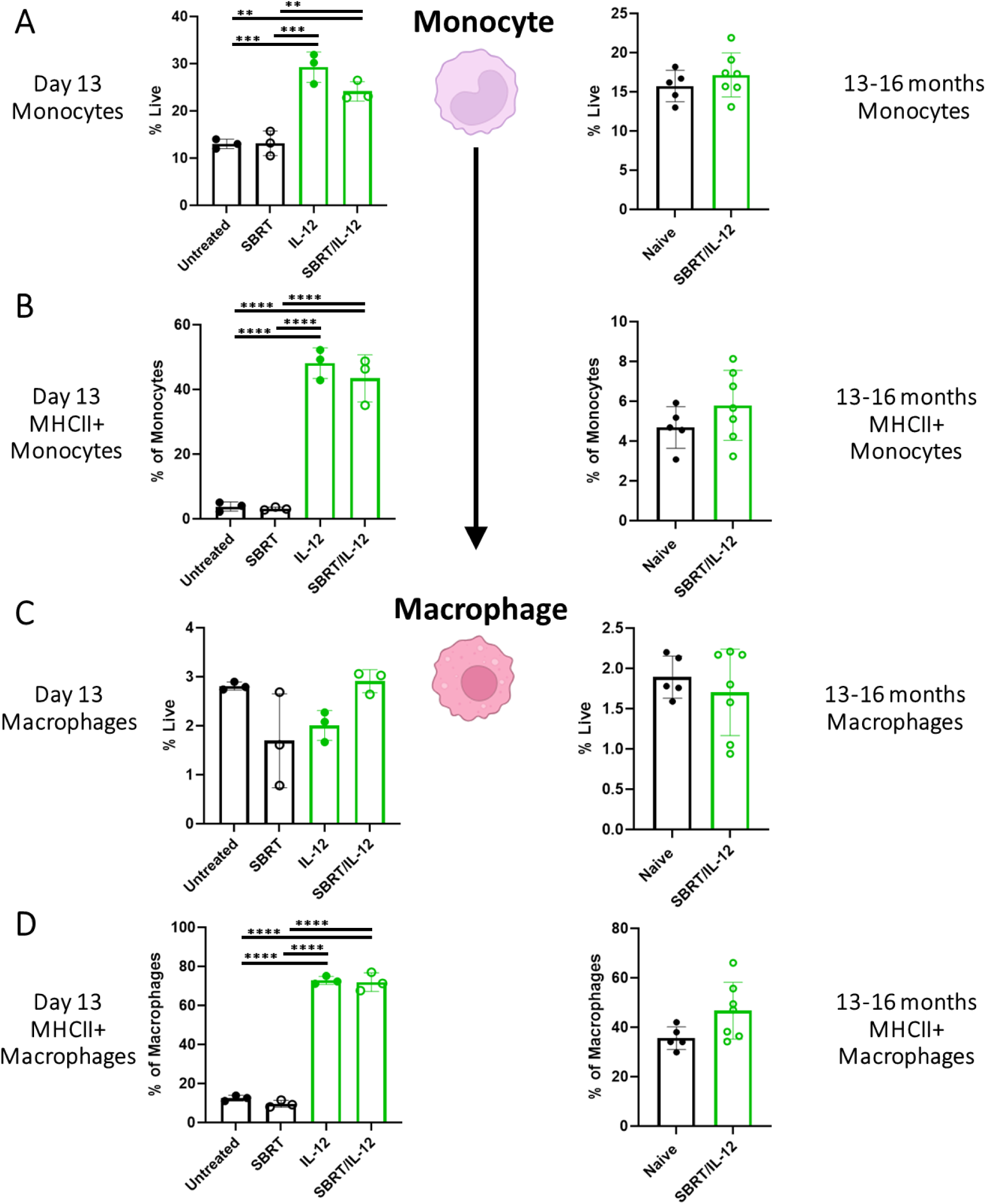
Bone marrow monocytes are temporarily increased and activated after SBRT/IL-12 therapy A and. **C** Percentage of Monocyte (CD45+, Ly6G-, Ly6C+) and macrophage (CD45+, Ly6G-, Ly6C-, F480+) as a frequency of live cells and **B and D** MHCII expression on monocytes and macrophages within the bone marrow 13 days and 13-16 months after tumor injection. n = 3-7 *p<0.05; **p<0.01; ***p<0.001; ****p<0.0001.

## 4. Discussion

In recent years, targeted therapies have become widely used in place of, or alongside systemic treatment due to reduced off-target effects [26]. Our laboratory utilizes a targeted combination of SBRT and IL-12mRNA that work together to overcome an aggressive form of PDAC [9]. SBRT induces immunogenic cell death (ICD), which signals dendritic cells (DCs) to stimulate and activate T cells [27, 28]. The factors released after SBRT and ICD can repolarize immunosuppressive myeloid cells in the tumor microenvironment (TME), but DCs fail to fully activate T cells within the milieu of immunosuppressive cytokines [29]. Local IL-12 enhances the antitumor response triggered by SBRT through stimulation of both innate and adaptive immune cells to overcome immune suppression and activate a potent antitumor response that completely eradicates PDAC tumors (**Figure 1**). Along with primary tumor eradication, local SBRT/IL-12 therapy provides systemic protection from liver metastases [8, 9]. This suggests that our local treatment, or the resultant immune response, could extend beyond the site of the tumor. Indeed, we observed changes in circulating lymphocytes and progenitor populations in the bone marrow within days of administering SBRT/IL-12. In light of the systemic alterations following local SBRT/IL-12 treatment, we conducted a detailed analysis of the hematopoietic populations in the bone marrow at multiple timepoints to determine the longevity of this response. Our findings highlight the importance of evaluating the long-term systemic effects of targeted therapies, as their impact could extend further than previously anticipated.

IL-12 plays an essential role in inducing T cells and NK cells to produce IFNγ, which is important for the robust antitumor T cell activation that SBRT/IL-12 therapy elicits [8, 9]. In line with this, we observed increased levels of IL-12 and IFNγ within the tumor shortly after treatment, confirming that both cytokines are rapidly elevated following therapy. However, IL-12 levels declined by 72 hours post-injection, indicating that the cytokine response is transient (**Figure 2**). These results are reassuring, as prolonged IL-12 exposure could lead to negative side effects [30, 31]. The IL-12 receptor is expressed mainly on NK cells and T cells; however, IL-12 stimulates these cells to produce IFNγ, which can directly interact with most cells in the body including lymphocytes, macrophages, and bone marrow progenitor cells [32–34]. Although our treatment is localized to the tumor, the pleiotropic effects of IL-12 and IFNγ could induce systemic changes if these cytokines disseminate throughout the blood. As such, lymphopenia was observed after SBRT/IL-12 therapy (**Figure 3**). Lymphopenia is a common result of conventional radiotherapy and chemoradiation therapy; however, SBRT has been shown to largely mitigate this effect [35, 36]. This suggests that IL-12 is likely responsible for the observed lymphopenia after therapy. This is supported by our data demonstrating that SBRT alone had little to no effect on both the peripheral blood and bone marrow metrics. In response to SBRT/IL-12 treatment, there is an influx of T cells in the tumor [9]. A mass migration of T cells from the blood to the inflammatory TME after therapy could explain the observed lymphopenia. Importantly, this effect was not permanent, as circulating levels returned to baseline 13-16 months after therapy (**Supplemental Figure 2**).

Our previous work has shown that SBRT/IL-12 therapy provides systemic protection against liver metastases, suggesting that other distal tissues, such as the bone marrow, could also be impacted. In this study, we confirmed that IL-12 levels in the bone marrow transiently increased 24 hours after pancreatic injection but returned to baseline by 72 hours, with levels 30-fold lower than in the tumor (**Figure 4**). Despite this small and brief increase, significant changes were observed in the bone marrow within 72 hours post-treatment. Previous studies have suggested that IFNγ plays a key role in mediating the effect of IL-12 on hematopoiesis by stimulating proliferation of HSCs [7, 33, 37, 38], however we found no evidence of increased IFNγ in the bone marrow after SBRT/IL-12 therapy. This suggests that IL-12 may act directly on stem cells, or IFNγ levels could be too low or rapidly consumed to detect. Further studies, including experiments with IFNγ knockout mice or assessing IFNγ at different timepoints, are needed to determine whether the observed changes are driven by IL-12 alone or in combination with IFNγ.

In order to effectively respond to a pathogenic challenge, immune cells are recruited to the site of infection. T cells have high proliferative potential, but myeloid cells, including macrophages, DCs, and monocytes, must be replenished from the bone marrow during an immune challenge in a process known as emergency hematopoiesis [11]. Our data suggest that emergency hematopoiesis may be induced by SBRT/IL-12 therapy (**Figure 5B**). This effect is predominantly due to the presence of IL-12, as evidenced by significant decreases in LT-HSCs in both IL-12 treatment groups. IL-12 can stimulate early hematopoietic stem cell progenitors which may lead to decreases in the overall population as they differentiate into immune cells [38]. Radiation could also play a role in the decrease of LT-HSCs, as previous studies have shown a noticeable shift in the bone marrow from hematopoietic red marrow to adipocyte rich yellow marrow even after indirect radiation [22, 39, 40]. Despite the potential for SBRT to alter the HSCs, our data did not reveal significant changes in the SBRT treated group, indicating that SBRT is not the driving factor for changes in the stem cell populations. In contrast to the LT-HSCs, the ST-HSC levels were not significantly different after therapy, but considering this is a transient population, it is not surprising that we saw no acute changes (**Figure 5D**). However, at the 13–16-month timepoint, we observed a decrease in ST-HSCs similar to the decrease in LT-HSCs, which is expected as there are fewer LT-HSCs to replenish the ST-HSC pool over time. Overall, these results illustrate the impact of local IL-12 on distal bone marrow stem cells.

In our murine model, we challenged the mice with tumors at an early timepoint and monitored their response shortly after treatment (day 13) as well as at a later stage (day 41), when the combination therapy group had achieved tumor clearance, and the mice were still considered young. We continued monitoring the mice into middle age (13-16 months) to assess the long-term effects of the therapy on the bone marrow microenvironment. During the natural aging process, the bone marrow undergoes changes involving the HSC populations, in which LT-HSCs expand, yet become increasingly dysfunctional [14]. Prior reports, including our own, have shown that bone marrow macrophages become dysfunctional with age, requiring HSCs to skew toward the myeloid lineage, indicated by an increase in CD41 expression [12, 41]. Similarly, we saw an increase in CD41 in our LT-HSC population and myeloid activation in our young mice soon after SBRT/IL-12 therapy (**Figure 5C**). While our results show that LT-HSCs expand with aging as expected, they remain decreased compared to controls at the 13 -16 month timepoint (**Figure 5B**). This would suggest these cells may be dysfunctional due to combination treatment. Together, these data hint at the possibility that our therapy may prematurely age the HSC population, resulting in an increased risk for clonal hematopoiesis and secondary myeloid neoplasms. Importantly, we challenged mice with tumor and treated them at a young age, whereas in a clinical context the median age of PDAC diagnosis is 71 [42]. Aged patients who receive SBRT/IL-12 therapy may be affected differently than younger patients since they would be expected to have a different bone marrow landscape. Future experiments could examine the effects of SBRT/IL-12 in an aged population of mice that better recapitulate the HSC populations seen in patients.

In alignment with increased CD41 expression on LT-HSCs, our findings indicate a dramatic acute increase in myelopoiesis after IL-12 therapy. MPP2 and MPP3 cell numbers were significantly higher at day 13 post tumor injection, however this was seen only in the IL-12 treated groups, indicating it is an IL-12 effect (**Figure 6A, Supplemental Figure 6A**). In the context of acute infection, IFNγ interacts with bone marrow stromal cells (BMSCs) to promote production of IL-6 which reduces expression of *Runx-1* and *Cebpα* to promote myeloid differentiation in HSCs [11, 37]. In our model, IL-12 may promote low and undetectable IFNγ production in the bone marrow, which could then increase myeloid differentiation. In clinical cases, high levels of IL-6 and increased myeloid cells in cancer patients are associated with poor prognosis [43]. In PDAC, the bone marrow serves as a source of myeloid-derived suppressor cells (MDSCs) that promote tumorigenesis and suppress antitumor immune cells within the tumor. PDAC tumors are characterized as having low numbers of tumor cells and a high proportion of immune and stromal cells [44], with a majority of these cells being immunosuppressive MDSCs from the bone marrow [45, 46]. This recruitment is due to tumor-derived factors (TDFs) that alter myelopoiesis in the bone marrow. Tumors secrete a variety of TDFs including IL-6, IL-10, vascular endothelial growth factor (VEGF), and granulocyte-monocyte colony-stimulating factor (GM-CSF) [43]. These cytokines play a role in suppressing the differentiation of immature myeloid cells and recruiting them to the tumor dampening the antitumor immunity. However, IL-12 has been shown to repolarize MDSCs to become functional antigen-presenting cells and recover macrophage functions [47, 48]. Repolarization involves an upregulation of MHCII expression on MDSCs and reduction of their suppressive effects on T cells [49]. We observed this in our studies as well, with an upregulation of MHCII expression on both monocytes and macrophages within the bone marrow 72 hours after the intratumoral IL-12 injection (**Figure 7**). This is in contrast to steady-state macrophages in the bone marrow, which do not express high levels of MHCII [25]. Our findings suggest that SBRT/IL-12 therapy can both repolarize the current suppressive cells within the TME and incite new production of activated myeloid cells that may assist in the antitumor response.

Our study has revealed minimal long-term effects induced by SBRT/IL-12 therapy, making it a promising choice for PDAC treatment. A major finding from this study was the prolonged decrease in LT-HSCs after tumor cure. Although this is not an ideal outcome, it is important to note that most standard cancer treatments, including radiotherapy and chemotherapy, also induce HSC senescence in the bone marrow [16]. This suggests that the effects observed from SBRT/IL-12 therapy are not unique, and remain within accepted limits. Moreover, in contrast to other treatments, our therapy can provide complete tumor cure. This demonstrates that the unprecedented benefits outweigh the associated risks, providing evidence that SBRT/IL-12 should continue to be developed for clinical treatment of PDAC.

