## Supplementary data for "Local delivery of SBRT and IL-12 to Murine PDAC Tumors Modulates Hematopoiesis"

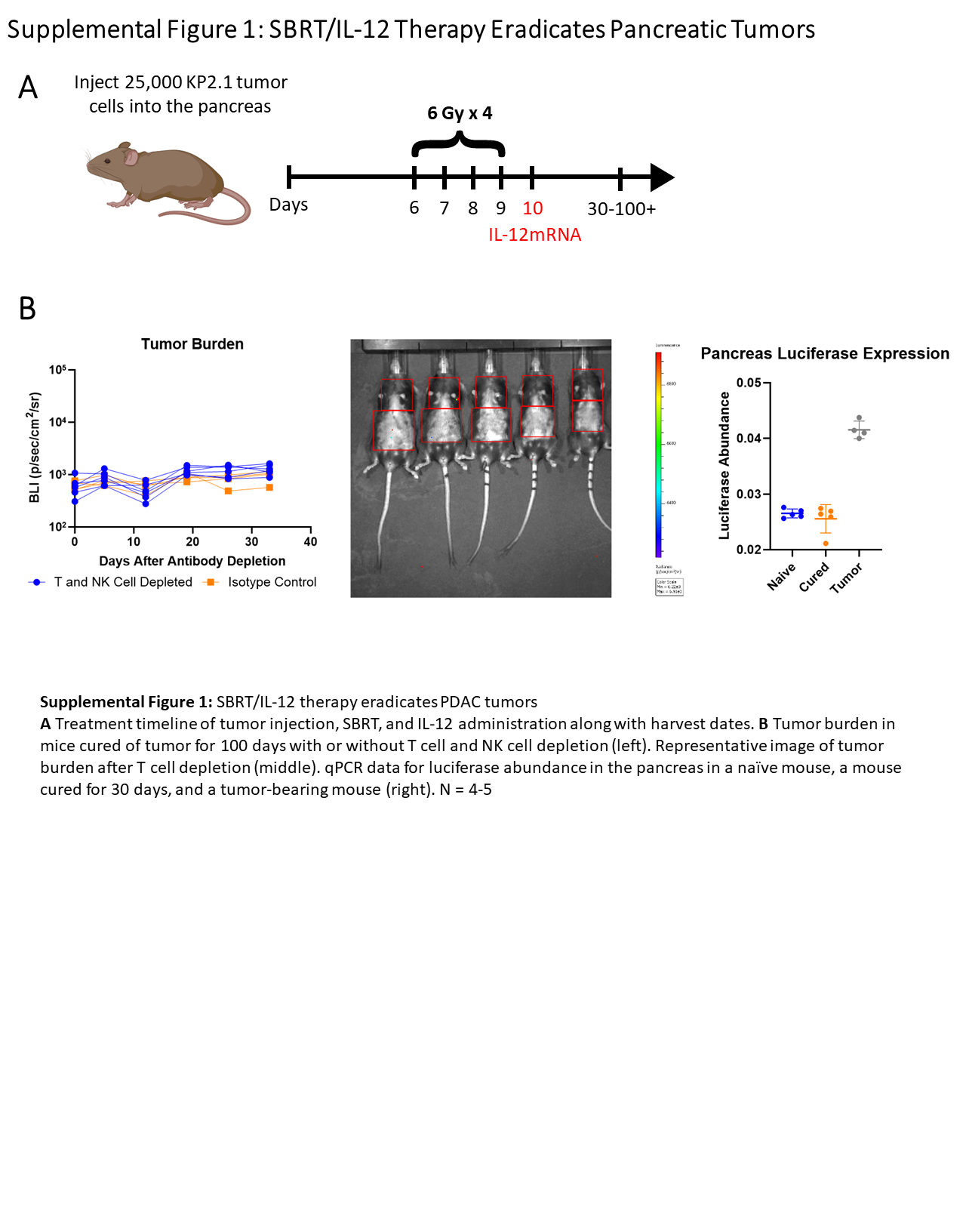

**Supplemental Figure 1: SBRT/IL-12 therapy eradicates PDAC tumors**

**A** Treatment timeline of tumor injection, SBRT, and IL-12 administration along with harvest dates. **B** Tumor burden in mice cured of tumor for 100 days with or without T cell and NK cell (left). Representative image of tumor burden after T cell depletion (middle). qPCR data for luciferase abundance in the pancreas of a naïve mouse, a mouse cured for 30 days, and the pancreatic tumor from a tumor-bearing mouse (right). n = 4-5.

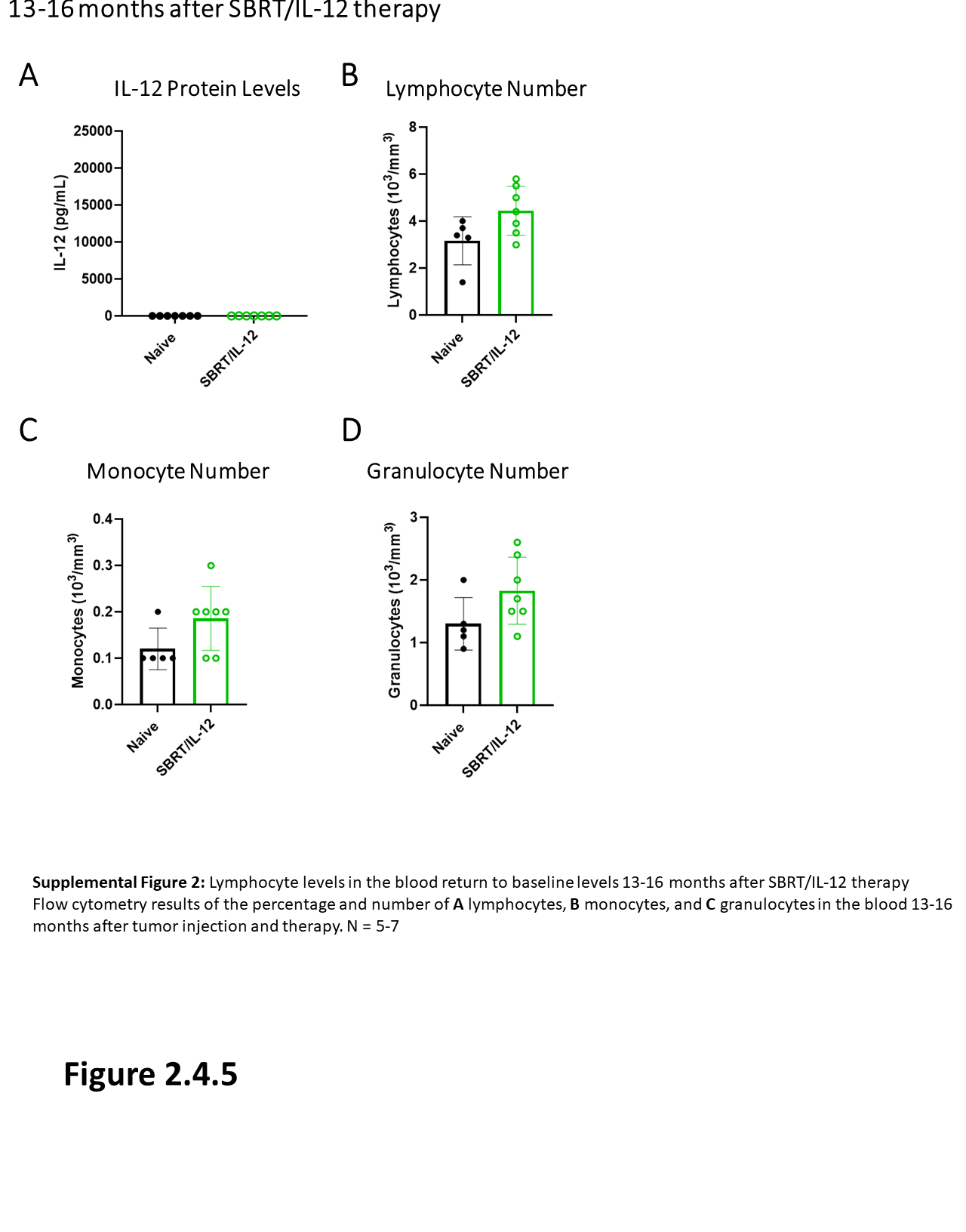

**Supplemental Figure 2: Lymphocyte levels in the blood return to baseline levels 13-16 months after SBRT/IL-12 therapy**

**A** ELISA data from blood serum harvested from mice 13-16 months after IL-12mRNA injection or naïve age-matched controls. Complete blood count (CBC) data showing the number of **B** lymphocytes, **C** monocytes, and **D** granulocytes in the blood 13-16 months after tumor injection and therapy. n = 5-7.

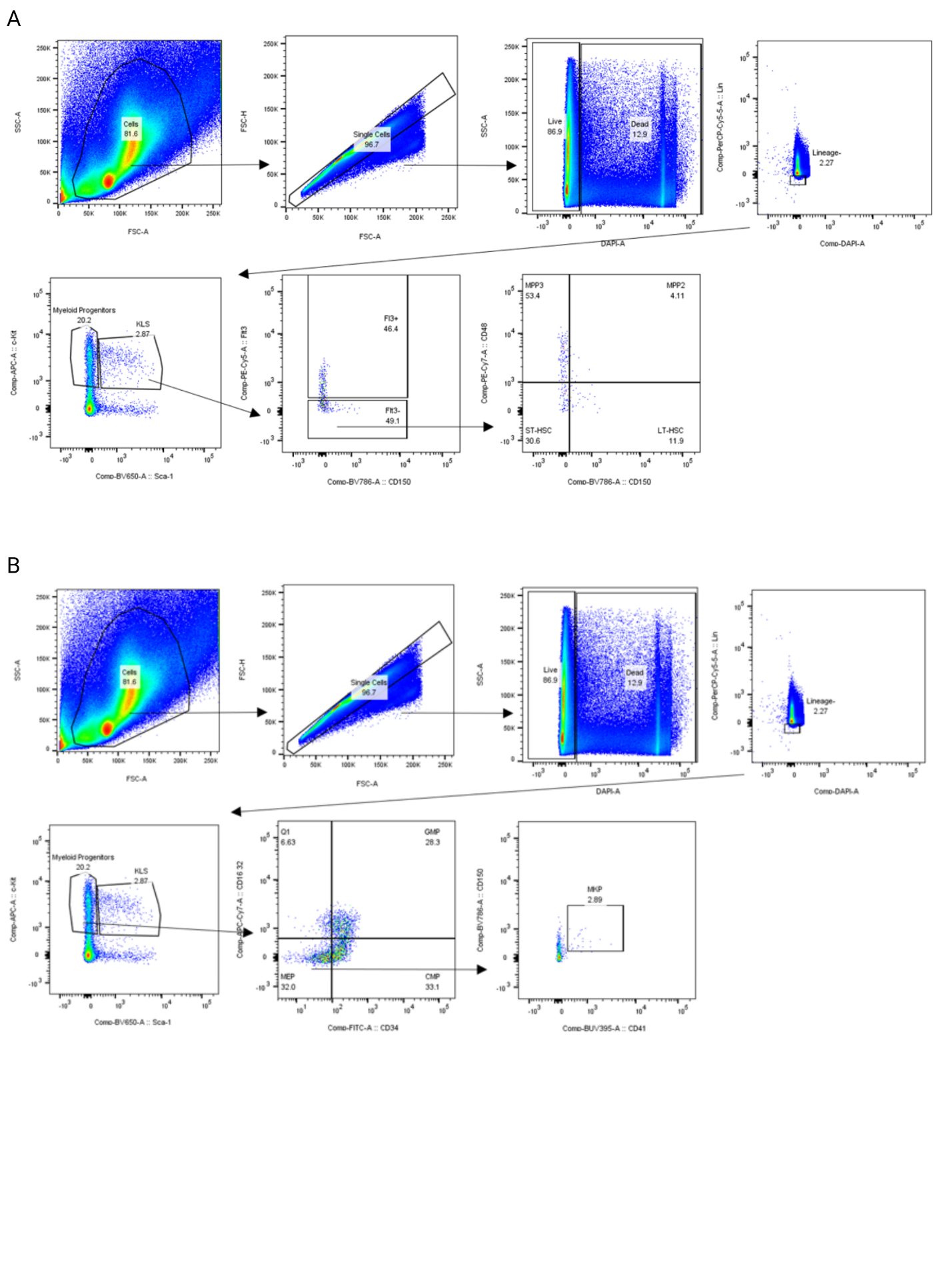

**Supplementary Figure 3: Gating strategy for identifying hematopoietic stem cells and their downstream progenitors.**

**A** Flow cytometry data showing the representative gating scheme for LT-HSCs (Lin-, c-Kit+, Sca-1+, Flt3-, CD48-, CD150+), ST-HSCs (Lin-, c-Kit+, Sca-1+, Flt3-, CD48-, CD150-), MPP2s (Lin-, c-Kit+, Sca-1+, Flt3-, CD48+, CD150+), and MPP3s (Lin-, c-Kit+, Sca-1+, Flt3-, CD48+, CD150-). **B** Flow cytometry gating scheme for CMP (Lin-, c-Kit+, Sca-1-, CD16/32-, CD34+), GMP (Lin-, c-Kit+, Sca-1-, CD16/32+, CD34+), MEP (Lin-, c-Kit+, Sca-1-, CD16/32-, CD34-), and MKP cells (Lin-, c-Kit+, Sca-1-, CD16/32-, CD34-, CD41+).

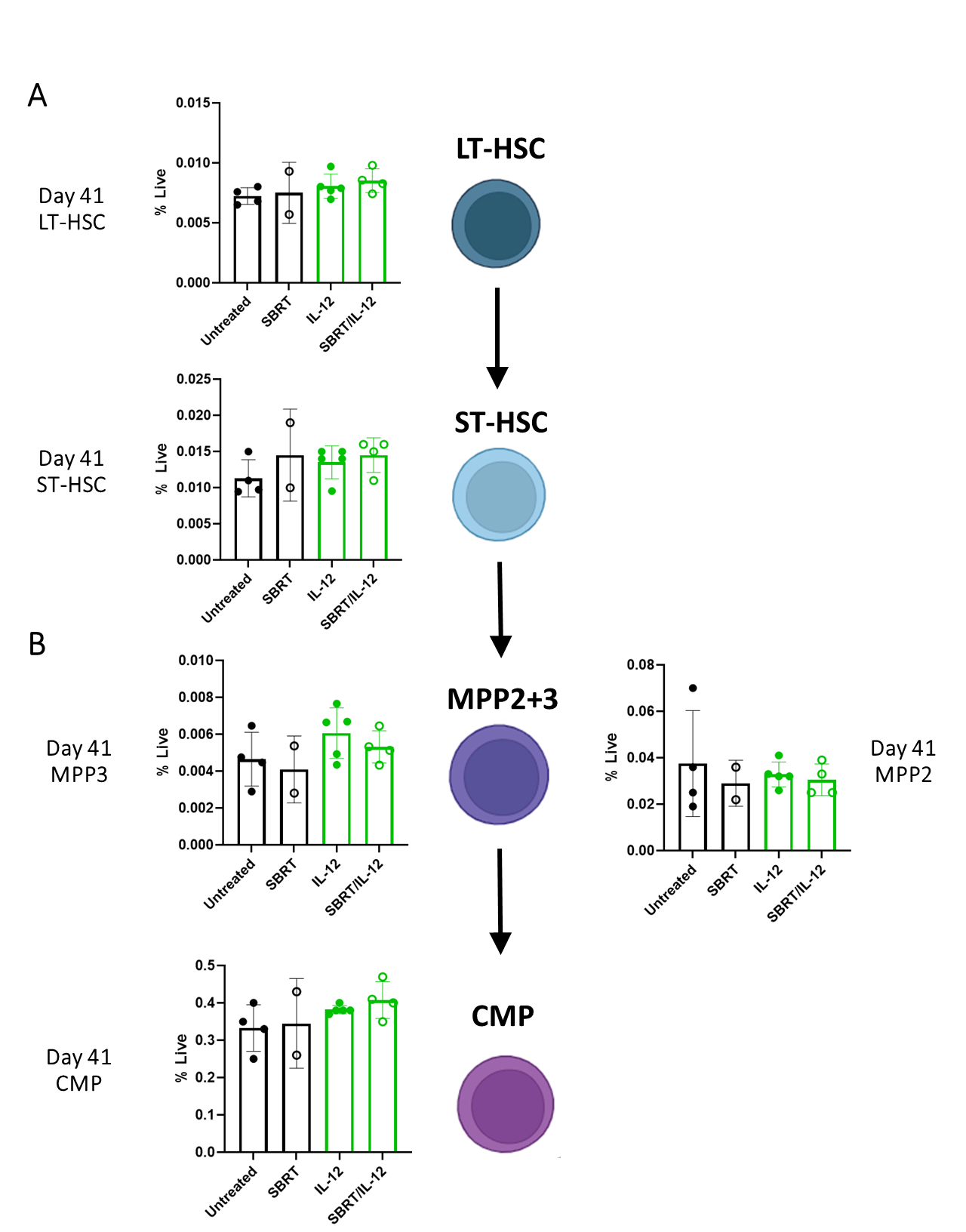

**Supplemental Figure 4: Progenitor cell numbers in the bone marrow normalize one month after therapy**

Flow cytometry data showing the percentage of **A** LT-HSCs, ST-HSCs, **B** MPP2 and MPP3 cells, and CMPs within the bone marrow 41 days after tumor injection. n = 2-4.

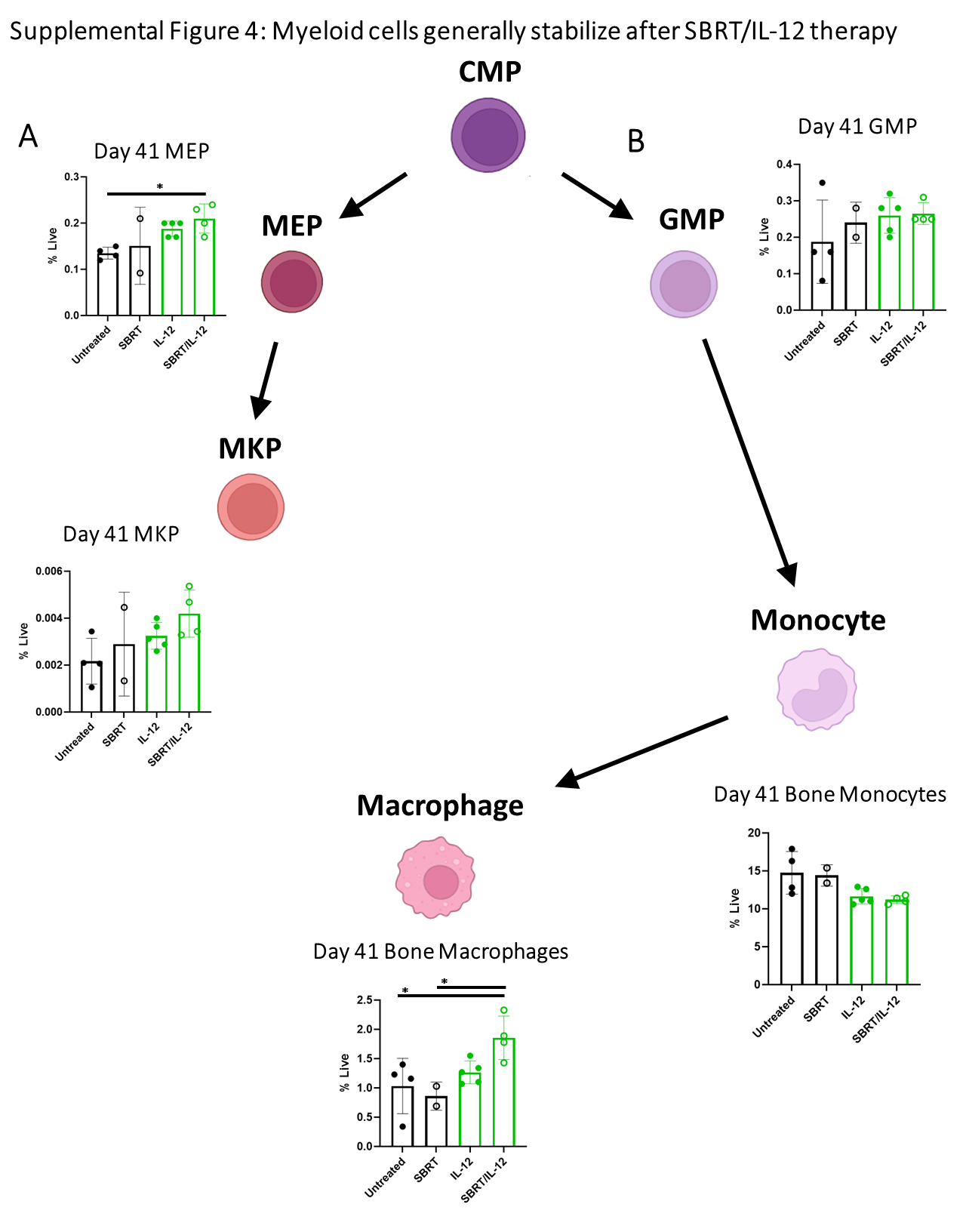
**Supplemental Figure 5: Myeloid cells generally stabilize after SBRT/IL-12 therapy**

**A** MEP and MKP levels in the bone marrow 41 days after tumor injection. **B** Percentages of GMPs, monocytes, and macrophages in the bone marrow 41 days post-tumor injection. Untreated and SBRT-treated mice received scRNA as a control. n = 2-4; *p<0.05.

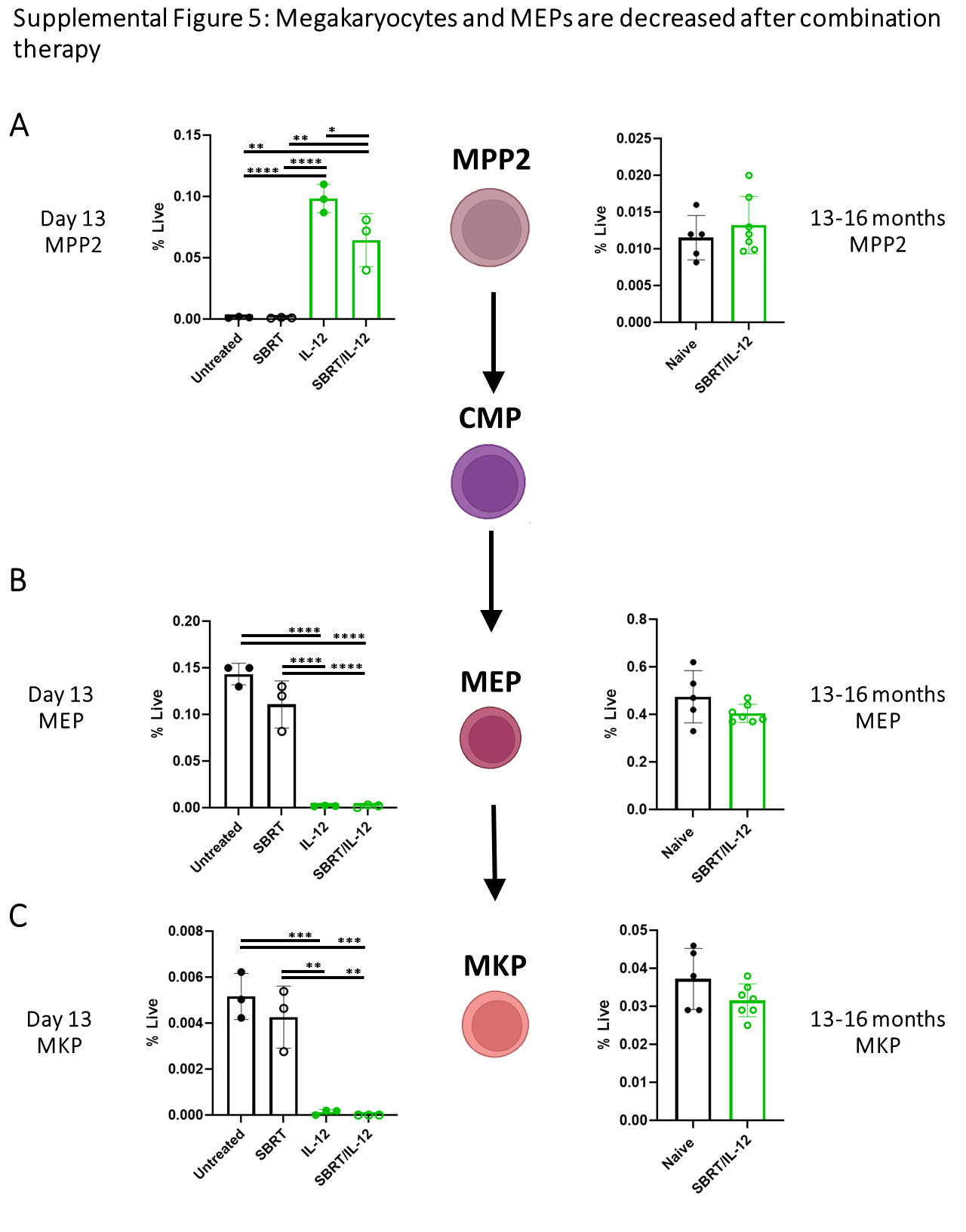

**Supplemental Figure 6: MKPs and MEPs are decreased after combination therapy**

Flow cytometry data showing the percentage of **A** MPP2, **B** MEP, and **C** MKP cells at day 13 and 13-16 months after tumor injection. One-way ANOVA with Tukey Correction. Untreated and SBRT-treated mice received scRNA as a control. n = 3-7; **p<0.01; ***p<0.001; ****p<0.0001.

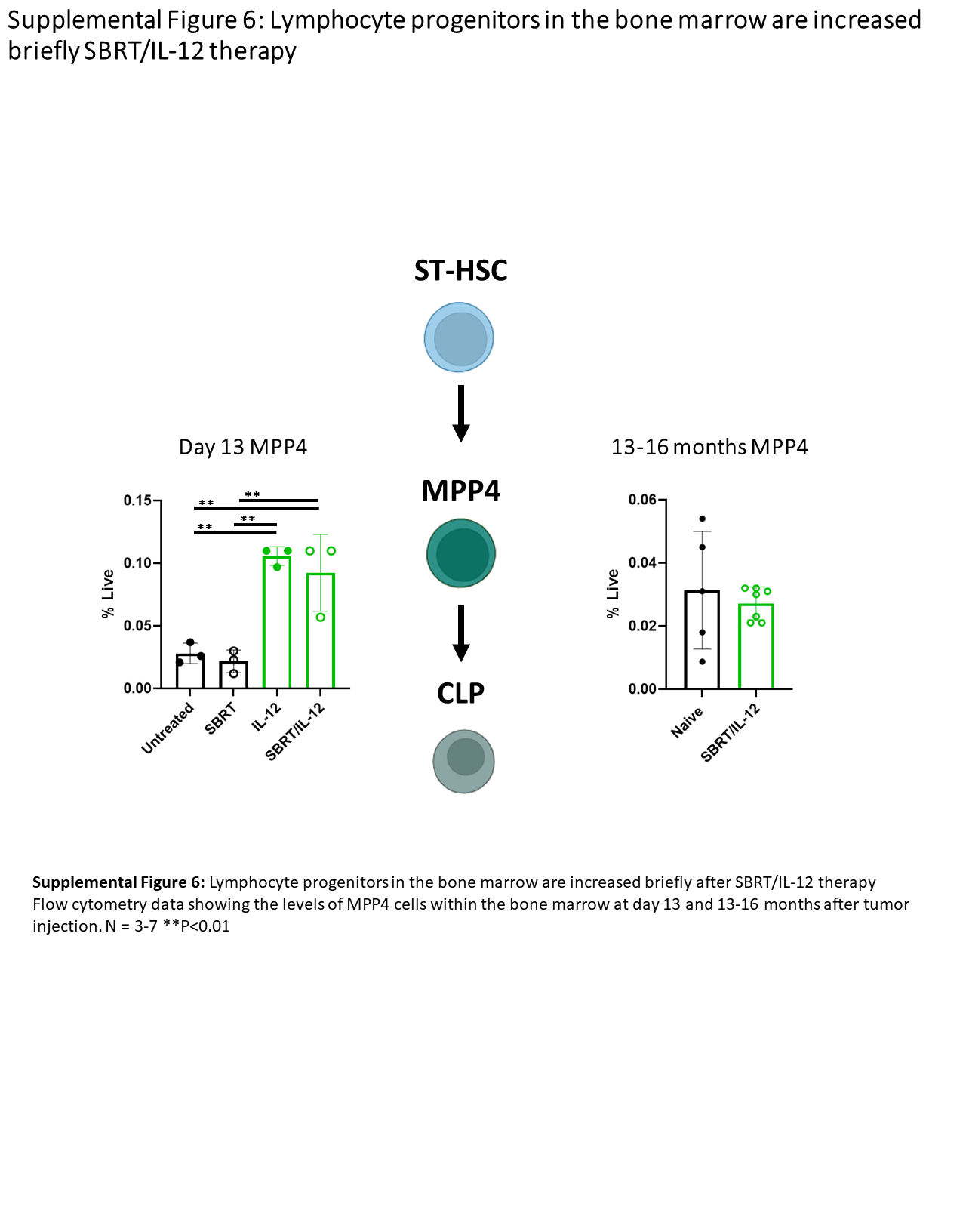

**Supplemental Figure 7: Lymphocyte progenitors in the bone marrow are increased briefly after SBRT/IL-12 therapy**

Flow cytometry data showing the levels of MPP4 (Lin-, c-Kit+, Sca-1+, Flt3+, CD48+, CD150-) cells within the bone marrow at day 13 and 13-16 months after tumor injection. Untreated and SBRT-treated mice received scRNA as a control. n = 3-7; **p<0.01.

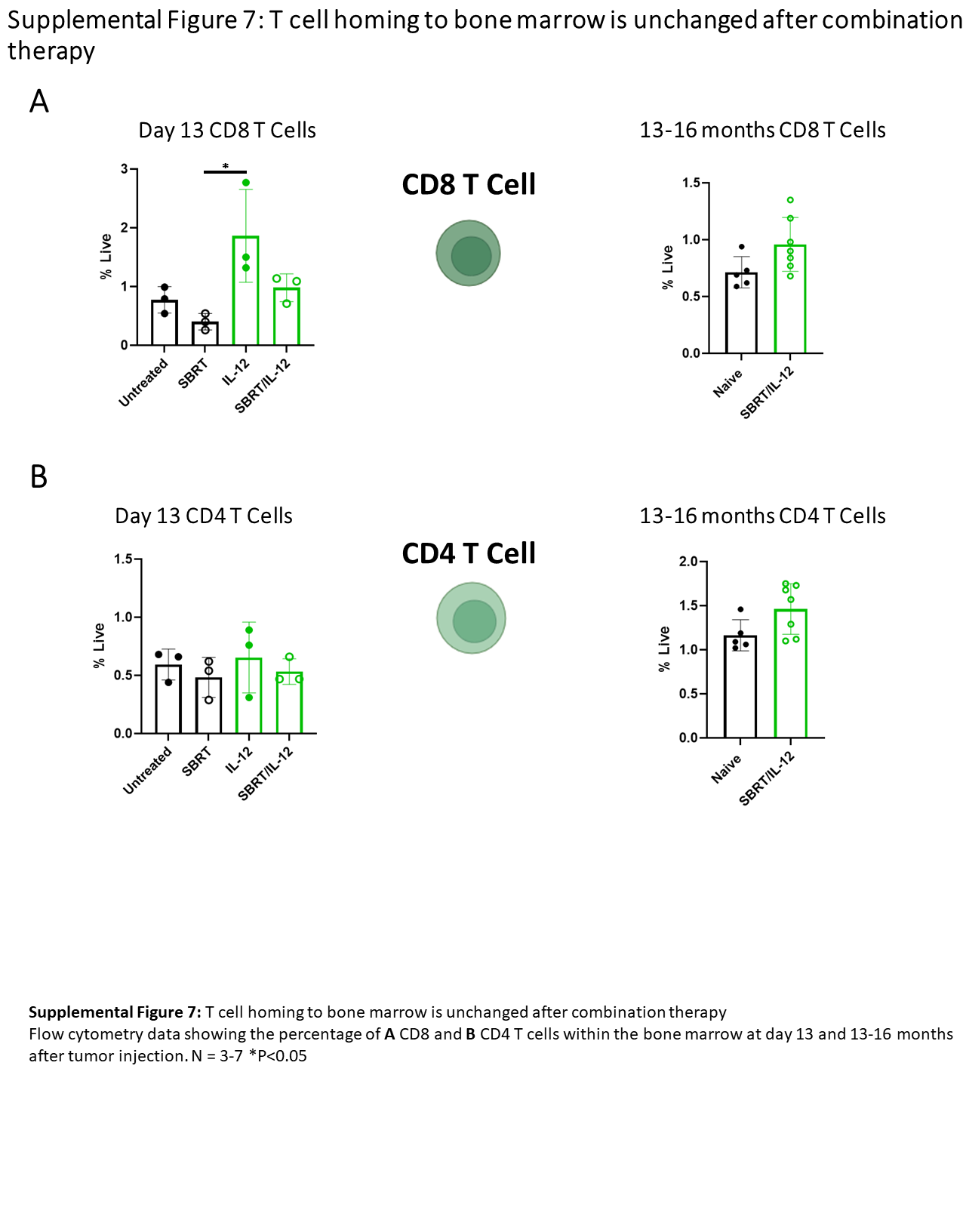

**Supplemental Figure 8: T cell homing to the bone marrow is unchanged after combination therapy**

Flow cytometry data showing the percentage of **A** CD8 (CD45+, CD11b-, CD3+, CD8+) and **B** CD4 (CD45+, CD11b-, CD3+, CD4+) T cells within the bone marrow at day 13 and 13-16 months after tumor injection. Untreated and SBRT-treated mice received scRNA as a control. n = 3-7; *p<0.05.

**Supplementary Table 1: Bone Marrow Cell Populations**

| **Cell Type** | **Flow Cytometry Markers** |
| --- | --- |
| **LT-HSC** | DAPI-, Lineage- (CD3, Ter119, Gr1, B220), Sca1+, cKit+, Flt3-, CD48- , CD150+ |
| **ST-HSC** | DAPI-, Lineage- (CD3, Ter119, Gr1, B220), Sca1+, cKit+, Flt3-, CD48- , CD150- |
| **MPP2** | DAPI-, Lineage- (CD3, Ter119, Gr1, B220), Sca1+, cKit+, Flt3-, CD48+ , CD150+ |
| **MPP3** | DAPI-, Lineage- (CD3, Ter119, Gr1, B220), Sca1+, cKit+, Flt3-, CD48+ , CD150- |
| **MPP4** | DAPI-, Lineage- (CD3, Ter119, Gr1, B220), Sca1+, cKit+, Flt3+, CD48+ , CD150- |
| **GMP** | DAPI-, Lineage- (CD3, Ter119, Gr1, B220), Sca1-, cKit+, CD34+, CD16/32+ |
| **CMP** | DAPI-, Lineage- (CD3, Ter119, Gr1, B220), Sca1-, cKit+, CD34+, CD16/32- |
| **MEP** | DAPI-, Lineage- (CD3, Ter119, Gr1, B220), Sca1-, cKit+, CD34-, CD16/32- |
| **MKP** | DAPI-, Lineage- (CD3, Ter119, Gr1, B220), Sca1-, cKit+, CD34-, CD16/32-, CD150+, CD41+ |
| **Monocytes** | DAPI-, Lineage- (CD3, Ter119, B220), CD45+, Ly6G-, Ly6C+, F480- |
| **Macrophages** | DAPI-, Lineage- (CD3, Ter119, B220), CD45+, Ly6G-, Ly6C-, F480+ |
